# Cellular and molecular mechanisms of frontal bone development in spotted gar (*Lepisosteus oculatus*)

**DOI:** 10.1101/2020.11.16.383802

**Authors:** Alyssa Enny, Andrew W. Thompson, Brett Racicot, Ingo Braasch, Tetsuya Nakamura

**Affiliations:** Department of Genetics, Rutgers the State University of New Jersey, Piscataway, NJ, 08854, USA; Department of Integrative Biology, Michigan State University, East Lansing, MI, 48824, USA; Program in Ecology, Evolution, and Behavior (EEB), Michigan State University, East Lansing, MI, 48824, USA

**Keywords:** spotted gar, frontal bone, dermal ossification, mesenchymal cell condensation

## Abstract

**Background:** The molecular mechanisms initiating vertebrate cranial dermal bone formation is a conundrum in evolutionary and developmental biology. Decades of studies have determined the developmental processes of cranial dermal bones in various vertebrate species, finding possible inducers of dermal bone. However, the evolutionarily derived characters of current experimental model organisms hinder investigations of the ancestral and conserved mechanisms of vertebrate cranial dermal bone induction. Thus, investigating such mechanisms with animals diverging at evolutionarily crucial phylogenetic nodes is imperative.

**Results:** We investigated the cellular and molecular foundations of skull frontal bone formation in the spotted gar *Lepisosteus oculatus*, a basally branching actinopterygian. Whole-mount bone and cartilage stainings and hematoxylin-eosin section stainings revealed that mesenchymal cell condensations in the frontal bone of spotted gar develop in close association with the underlying cartilage. We also identified novel aspects of frontal bone formation: Upregulation of F-actin and plasma membrane in condensing cells, and extension of podia from osteoblasts to the frontal bone, which may be responsible for bone mineral transport.

**Conclusion:** This study highlights the process of frontal bone formation with dynamic architectural changes of mesenchymal cells in spotted gar, illuminating supposedly ancestral and likely conserved developmental mechanisms of skull bone formation among vertebrates.

## 1 | INTRODUCTION

The patterning and growth of cranial dermal bones in vertebrates are remarkably diverse and often reflect functionally different demands in various habitats^1,2^. The functional importance and complexity of cranial dermal bones in vertebrates are illustrated by the skull roof, which primarily protects the brain and sensory organs. More than 450 million years ago, large bony plates covered the primitive fish cranium for protection against predators. These plates, known as the macromeric condition, originated at the dorsal and ventral cranium in groups of ancient jawless fishes such as Arandaspida, Heterostraci, and Osteostraci^3–5^. Likewise, jawed fishes of the group Placodermi possessed large bony plates that covered the head and the anterior body trunk. The evolutionary relationship of Placodermi’s bony plates that have a peculiar size and shape to the skull bones of other fishes is currently under investigation^6–8^. In the course of vertebrate evolution, a myriad of diverse dermal skull roof patterns were converged to the archetype represented by extant actinopterygians and sarcopterygians^9^. Despite centuries of anatomical studies, however, the genetic underpinnings of morphological diversity in the skull roof remain poorly understood^10–15^.

The evolutionary diversity of the skull roof could be revealed by identifying where and how ossification for each bone initiates in the embryonic head. During embryonic skull roof formation, mesenchymal cell condensations, which later differentiate to osteoblasts, form between the epidermis and endocranial cartilage, giving rise to the skull roof. A previous study in cichlid fishes used transmission electron microscopy to show that spindle-shaped cells among ubiquitously distributed mesenchymal cells congregate with gap junctions and form mesenchymal cell condensations^16^. Gap junctions are indispensable for the ossification process, presumably allowing diffusion of ions or small signaling molecules between condensing cells to induce osteoblast differentiation for skull roof ^17–20^. The cranial dermal bones ossify exclusively with osteoblasts, thus lacking the chondrocytes present in endochondral ossification, and undergo what is referred to as intramembranous ossification^21–23^.

The molecular mechanisms that induce mesenchymal condensations for cranial dermal bone ossification remain enigmatic^24^. Historical studies identified their close localization with lateral line neuromasts^25–28^, endocranial cartilage^2,23,29^, and epithelial tissues^23,24,30^. However, there is no decisive evidence to conclude that these tissues induce mesenchymal cell condensations for cranial dermal bones. In early studies, close association of cranial dermal bones with lateral lines in extinct and extant taxa suggests that chemical signals or cell migrations from neuromasts stimulate the formation of mesenchymal condensations during cranial dermal bone development^31–33^. However, other studies have been reluctant to support this lateral line hypothesis, recognizing close topology between the cranial dermal bones and lateral lines as a developmental coincidence^24^. As such, laser removal of neuromasts from the lateral line housed in the infraorbital bones does not affect overall shape of the infraorbital bones^34^.

Recent mouse studies have shed light on the functional necessity of epithelial cells in initiation of skull roof formation. Secreted signaling proteins fibroblast growth factors 8 and 10 (Fgf8 and Fgf10) are expressed in facial ectodermal cells (epithelial cells)^35,36^, and their receptor Fgfr2 is expressed in underlying mesenchymal cells^37^. Conditional deletion of *Fgfr2* in the mesenchymal cell layer by *Dermo1:cre* results in malformed skull roof bones^37^, implying that Fgf8, Fgf10, and presumably other Fgf ligands regulate skull roof ossification processes via diffusion from the ectoderm to mesenchymal cells. Along with Fgf ligands, various Wnt ligands are expressed in the facial ectoderm^38^. Deletion of *Wntless* (*Wls*), a Wnt ligand transporter^38^, in an ectoderm-specific manner causes arrest of osteoblast differentiation but has no effect on osteoblast proliferation in mouse. This phenotype implies that Wnt ligands expressed in the ectoderm diffuse into the mesenchymal cell layer with Wls and regulate cranial dermal bone formation^38^.

The skull roof develops via dermal ossification, in which cranial mesenchymal cells directly differentiate into osteoblasts^39^. Despite a significant number of studies on skull roof development using diverse model organisms with evolutionarily derived features^40^, the ancestral conditions of skull roof developmental mechanisms are relatively unexplored. Prior studies have used whole-mount bone staining and sectioning to describe the developmental process of skull roof formation in basal actinopterygian fishes including the holosteans gar (Lepisosteidae) and bowfin (*Amia calva*), which hold a phylogenetically prominent position in vertebrate evolution as an outgroup to teleost fishes^31,32^. However, investigation of the structure and developmental mechanisms of the skull roof in these basal actinopterygians at higher resolution is critical to better elucidate the ancestral and shared mechanisms of this process.

Thus, we used the holostean spotted gar (*Lepisosteus oculatus*), which retains a genome structure that is more comparable to the genomes of sarcopterygian vertebrates including humans than to those of teleost fishes^41^. The genome of the spotted gar has not undergone the teleost-specific genome duplication as developmental models like zebrafish and medaka did. In addition, the very slow morphological and molecular evolutionary rates of gar make it an important model to infer ancestral molecular mechanisms responsible for skull roof formation. We conducted single-cell analysis of frontal bone formation, one of skull roof bones, in spotted gars with fine hematoxylin and eosin (HE) staining, 3D reconstruction, and cytoskeleton and plasma membrane staining. The results redefine the processes involved in frontal bone development and highlight novel mesenchymal cell behavior during frontal bone development. This newly obtained knowledge will serve as a basis for future molecular and genetic studies of basal actinopterygian skull development, illuminating supposedly ancestral and shared developmental mechanisms of skull formation among the diversity of vertebrates.

## 2 | Results

### 2.1 | Development of the skull bones in spotted gar

To investigate developmental mechanisms of the gar skull roof in detail, we first re-examined developmental processes of the skull bones including the frontal bone. To accomplish this, we used acid-free bone and cartilage staining, which stains calcification with higher sensitivity than canonical bone staining methods^42^. Until 20 mm total length (TL), we did not find any evidence for the frontal bone or other skull roof bone development (data not shown). At 21 mm total length (TL), premaxillary, maxillary, and ectopterygoid bones (dermal bones) were stained by alizarin red S (Figure 1A, B). The parasphenoid, which lies at the roof of the mouth, extended to the anterior extremity of the head. Anterior to the pectoral fin, a series of pectoral girdle bones (extrascapular, supracleithrum, and cleithrum) extended from the posterior skull to the ventral edge of the body (Figure 1B). A transparent thickened anlage for the frontal bone was observed dorsal to the eye under a stereo-type microscope, yet alizarin red S staining for this anlage was not observed (Figure 1B’). This is consistent with previously defined developmental stages of gar skull bones^31,32,43^.

**Figure 1.**
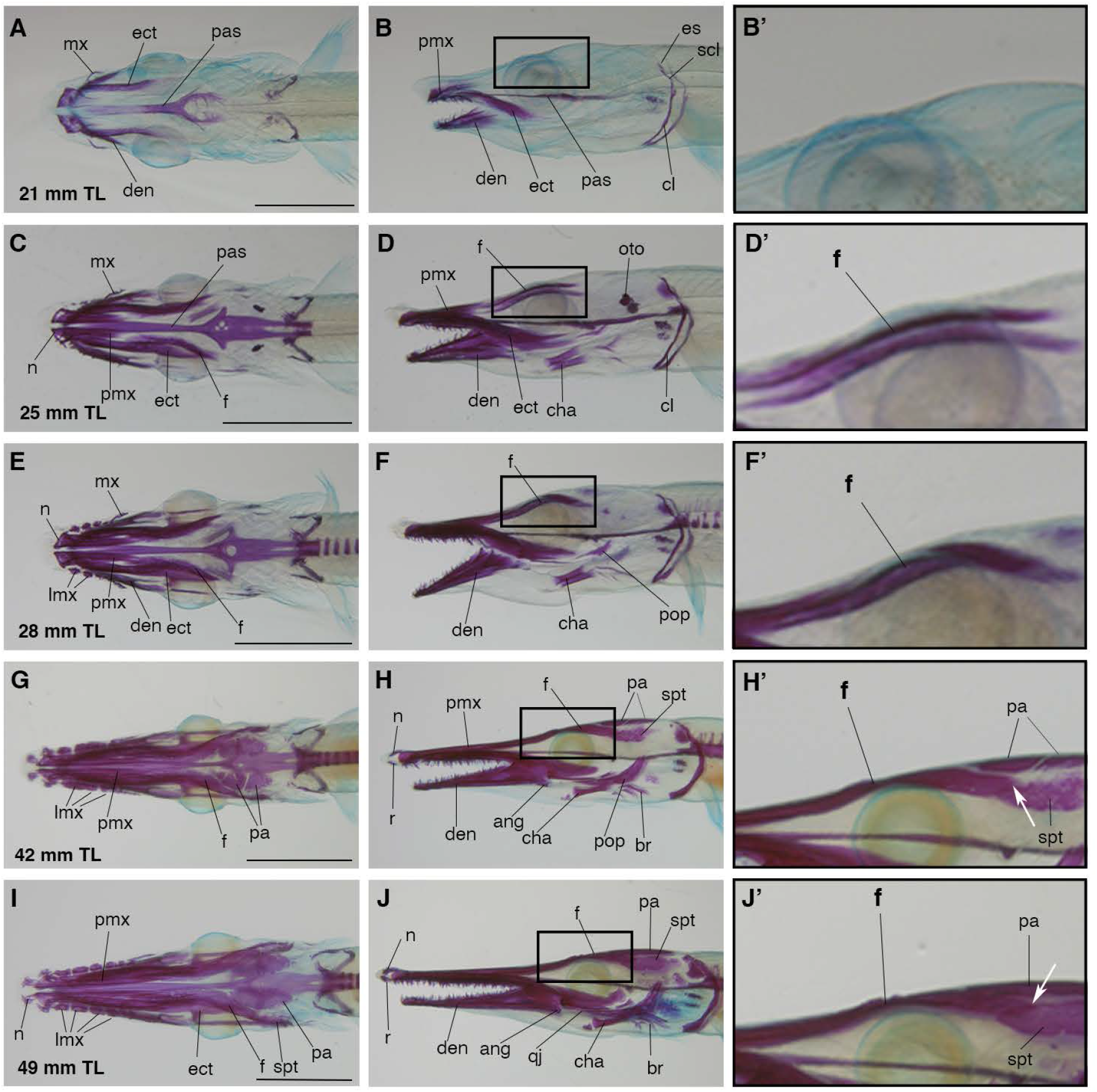
The developmental process of gar skull bones. A, C, E, G, and I; the dorsal views of acid-free bone and cartilage staining on 21 mm (A), 25 mm (C), 28 mm (E), 42 mm (G), and 49 mm (I) TL stage gar juveniles. B, D, F, H, and J; the lateral views of the same juveniles as A, C, E, G, and I. B’, D’, F’, H’, and J’; the magnification of the frontal bone domains in B, D, F, H, and J. At 21 mm TL stage (A, B, and B’), there is no evidence of ossification of the frontal bone (f). The ossification of jaw bones, including premaxilla (pmx), dentary (den), and ectopterygoid (ect) bones are discerned. Parasphenoid (pas) starts to ossify. The cleithrum (cl), supracleithrum (scl), and extrascapular (es) develop at the posterior to the head. At 25 mm TL stage (C, D, and D’), the frontal bone (f) began to ossify at dorsal to the eye. The premaxillary, dentary, and ectopterygoid bones are further ossified. The posterior end of premaxillary and the anterior end of the frontal bone has a gap between them (C). At 28 mm TL stage (E, F, and F’), the frontal bone extends to the anterior direction. The posterior end of the premaxillary bone and the anterior end of the frontal bone is still separated by a small gap. Several lacromaxillary (lmx) bones are discerned at the lateral to the ectopterygoid bone. At 42 mm TL stage (G, H, and H’), the frontal bone continues to grow anteriorly and posteriorly. The posterior end of the frontal bone is located at the lateral to the developing parietal bone (pa) (G and H’, white arrow). Parietal bones develop from the separated ossification centers. The supratemporal (spt) bone is observed posterior to the frontal bone (H’). At 49 mm TL stage (I, J, and J’), the premaxillary and frontal bones cover the anterior medial part of the skull. The separated parietal bones fuse with each other and expand, covering the top of the skull. White arrows in H’ and J’ indicate the posterior extremity of the frontal bone. Ang; angular, br; branchiostegals, cha; anterior ceratohyal, cl; ceithrum, den; dentary, ect; ectopterygoid, es; extrascapular, f; frontal, lmx; lacrimomaxilla, mx; maxilla, n; nasal, oto; otolith, pa; parietal, pas; parasphenoid, pmx; premaxilla, pop; preopercle, qj; quadoratojugal, r; rostral bone, sct; supracleithrum, spt; supratemporal, and pt; posttemporal. Anatomical terminology is based on Grande, L., 2010^74^. Scale bars are 5 mm.

At 25 mm TL, cranial dermal bones, including premaxillary, dentary, and ectopteryogoid bones, are further ossified (Figure 1C, D). The parasphenoid posteriorly reached the vertebral column and was more ossified at the 25 mm TL stage compared to the 21 mm TL stage (Figure 1C). At 25 mm TL, the frontal bone was clearly stained by alzarin red S (Figure 1D). The frontal bone developed dorsally to the supraorbital cartilage, a dorsal endocranial cartilage encasing the eye (Figure 1D’). The frontal bone stretched along the anteroposterior axis, and the anterior extremity was located close to the posterior end of the premaxillary bone. Alzarin red S staining for the frontal bone was strongest dorsal to the eye, and staining became weaker as it extended in anterior and posterior directions, indicating that frontal bone ossification starts with mesenchymal cells located dorsally to the eye.

At 28 mm TL, lacromaxillary bones were aligned lateral to the ectopterygoid bone (Figure 1E). The premaxillary bone extended posteriorly and approached the anterior end of the frontal bone (Figure 1E). The frontal bone slightly extended more to the anterior and posterior direction compared to 25 mm TL (Figure 1E, F). At this stage, the frontal bone became wider around the eye region (Figure 1E, F’). The preopercular bone started to ossify posteroventral to the eye.

At 42 mm TL, frontal and premaxillary bones further developed along the anteroposterior axis and could be distinguished under a stereomicroscope (Figure 1G). Posterior to the frontal bone, the parietal bone developed from multiple independent ossification centers (Figure 1G, H, H’). Frontal and parietal bones were separated by a distinct gap between them. Also, the supratemporal bone developed posterior to the frontal bone and lateral to the parietal bone (Figure 1H, H’). The posterior half of the frontal bone became wider than the anterior half.

Finally, at 49 mm TL, the frontal bone fully developed, with its posterior extremity between the parietal and supratemporal bones (Figure 1I, J’). The originally separated parietal bones fused with each other and constructed a large, flat osseous plate that dorsally covered the head (Figure 1I, J’). Supratemporal and preopercular bones ossified more compared to the previous 42 mm TL stage (Figure 1J, J’). Posterior to the angular bone, the quadratojugal bone started to develop (Figure 1J). For all stage juveniles, three individuals were investigated.

Overall, the observed acid-free bone and cartilage staining in spotted gar was consistent with previously published data^32,43^ and serves as a precise developmental staging series for subsequent fine analysis of gar skull roof development.

### 2.2 | 3D reconstruction of topology of the frontal bone, endocranial cartilage, and neuromasts

Prior studies in actinopterygians and sarcopterygians suggest that the frontal bone develops in the mesenchymal layer in close proximity to the underlying endocranial cartilage, epithelial cells, and lateral line neuromasts, positing these tissues as inducers of mesenchymal cell condensations^33,44,45^. However, the functional roles of these tissues in the initiation of mesenchymal condensations remain poorly understood. To precisely characterize topology of the frontal bone anlage and its possible inducers in spotted gar, we reconstructed the 3D morphology of the frontal bone anlage, endocranial cartilage, and neuromasts from HE-stained paraffin sections using Amira 3D analysis software.

At 14 mm TL, there was no indication of mesenchymal cell condensations for the frontal bone (Supplementary Figure 1A, A’). Supraorbital cartilage and neuromasts were observed dorsal to the eye.

At 17 mm TL, fragmented frontal bone anlagen were observed dorsal to the supraorbital cartilage (Figure 2A). This indicates that formation of anlagen for the frontal bone actually occurred earlier than the ossification stage identified by bone staining (Figure 1C, 25 mm TL stage, see above). Some of these anlagen were right under the neuromasts in the supraorbital lateral line (Figure 2A’). This result is consistent with previous teleost and non-teleost actinopterygian studies asserting that mesenchymal cell condensations for the frontal bone develop in close proximity to neuromasts in the supraorbital lateral line^2,29,33^. However, the other anlagen did not develop right below the neuromasts (Figure 2A’). At a more lateral position of the head, fragmented mesenchymal cell condensations also developed close to the neuromasts, yet these condensations were not right under the neuromasts either (Supplementary Figure 1B, B’).

**Figure 2.**
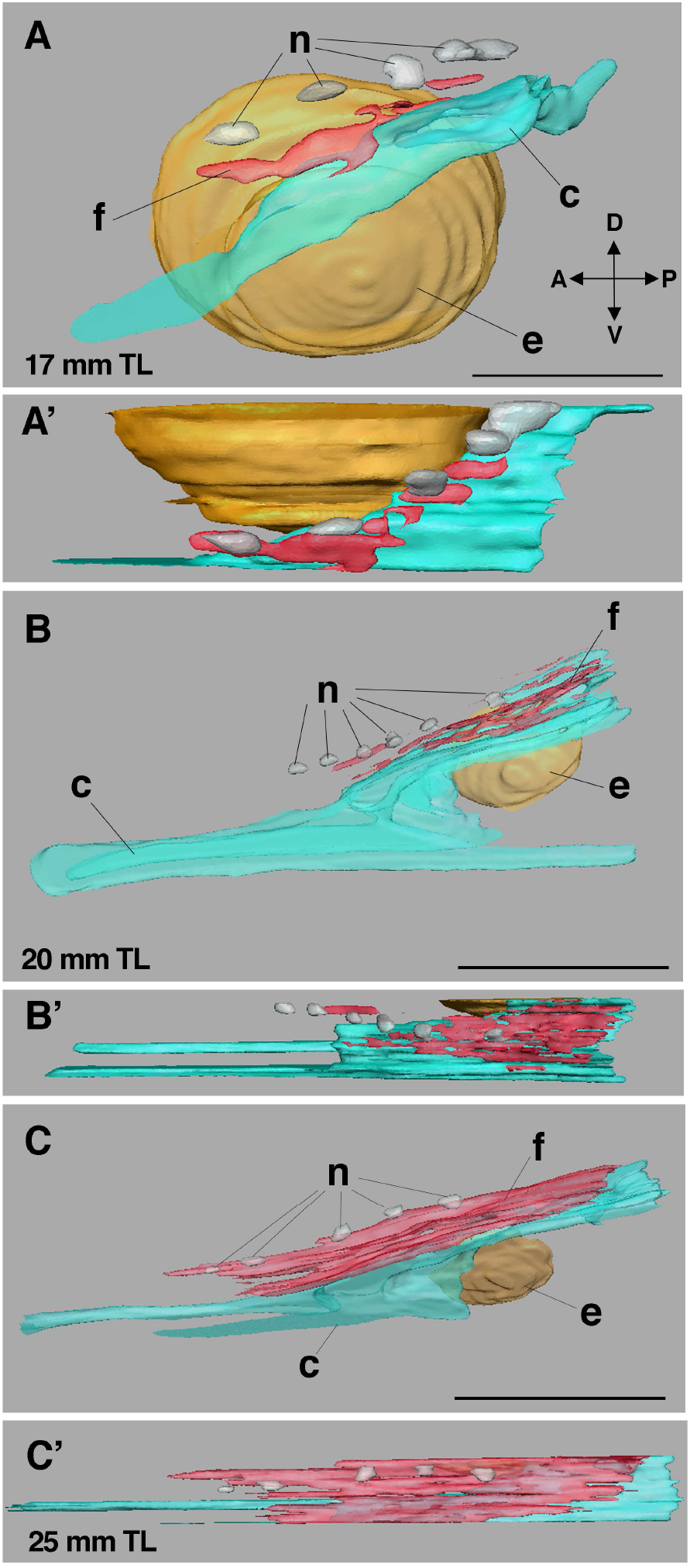
Topological relationships among the neuromasts, endocranial cartilage, and frontal bone. A, B, and C; the dorsomedial views of 3D reconstruction of the eye regions of gar skull at 17 mm (A), 20 mm (B), and 25 mm (C) TL stages. A’, B’, and C’; the dorsal views of the same juveniles of A, B, and C. At 17 mm TL stage (A and A’), the frontal bone anlage (f) is located dorsally to the endocranial cartilage (c). Although the neuromasts and the frontal bone anlage develop in a close proximity, their positions do not completely match (A’). At 20 mm TL stage, the frontal bone anlage anteriorly extends and cover the supraorbital cartilage above the eye (B and B’). The dorsal view of the developing neuromasts and frontal bone showed that the frontal bones do not develop exactly at the ventral to the neuromasts. At 25 mm TL stage, the frontal bone expands anteriorly and posteriorly (C and C’). While the frontal bone forms under the neuromasts, it also extends laterally without neuromasts (C’). The scale bars are 0.2 mm. n; neuromasts, e; eye, c; endocranial cartilage, and f; frontal bone.

At 20 mm TL, frontal bone anlagen fused with each other and extended along the anteroposterior axis. These anlagen were located dorsal to the supraorbital cartilage, which encases the eye (Figure 2B, B’). Although the frontal bone anlage developed in close proximity to the neuromasts, positions of the neuromasts and frontal bone anlage did not precisely match (Fig. 2B’). In addition, the frontal bone anlage expanded laterally and posteriorly, independent of neuromasts (Figure 2B’).

At 25 mm TL, the frontal bone anlage became a thin and flat osseous plate that dorsally covered the supraorbital cartilage. The frontal bone was located under the neuromasts, although the edge of the bone extended laterally without neuromasts. Also, the bone significantly extended in anterior and posterior directions, which did not seem to correlate with neuromast positions (Figure 2C, C’). For all stage juveniles, two individuals were investigated.

Overall, these results support the assertion that the proximity of neuromasts and ossification centers for cranial dermal bones is a developmental coincidence and not causally linked^24^, at least in spotted gar.

### 2.3 | Cellular basis of frontal bone development

To further scrutinize the process of frontal bone development, HE-stained paraffin sections of gar juveniles from 14 mm TL to 25 mm TL were observed at single-cell resolution (Figure 3).

**Figure 3.**
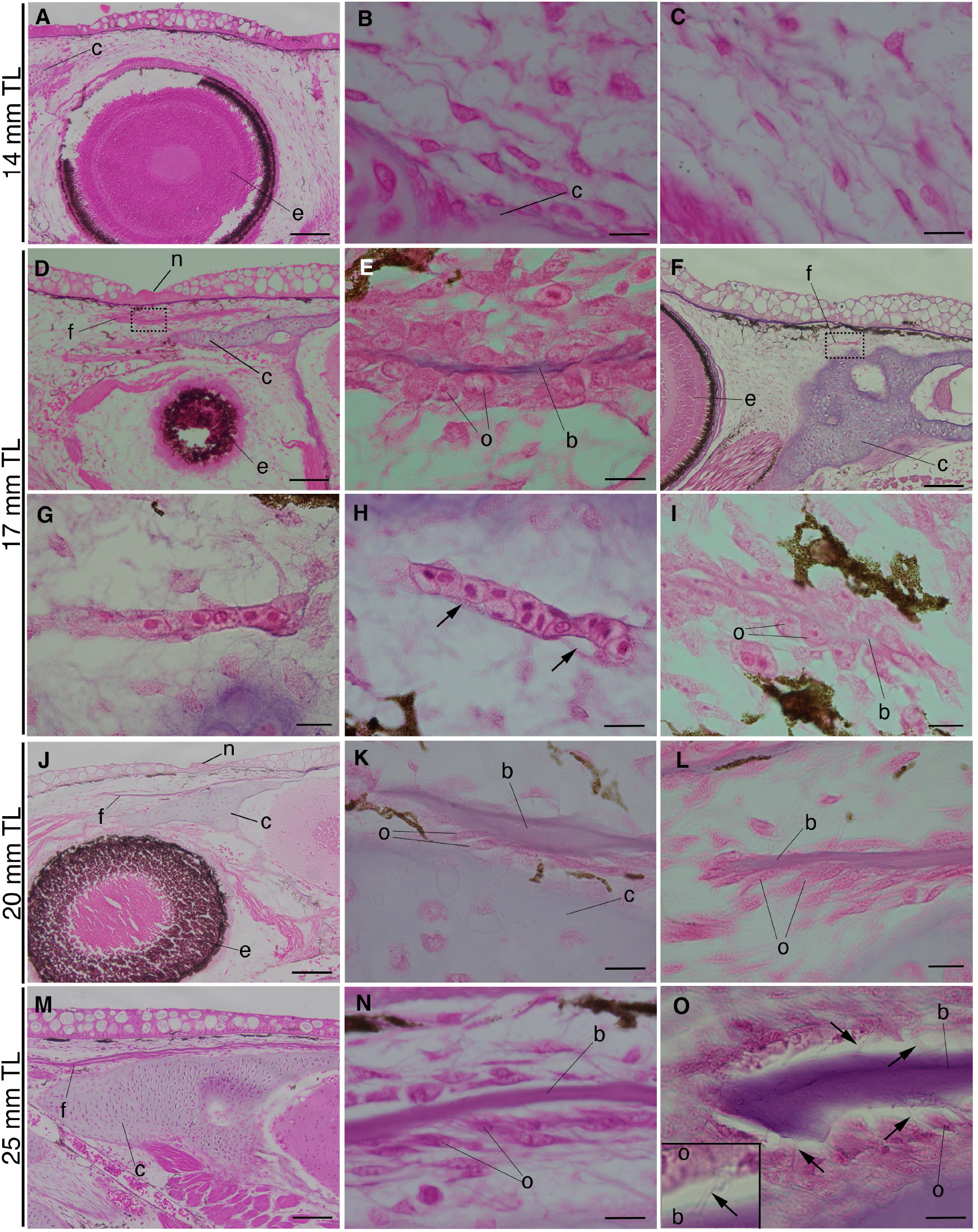
Developmental process of the frontal bone at single-cell resolution. A-C; 14 mm TL, D-I; 17 mm TL, J-L; 20 mm TL, and M-O; 25 mm TL stages. A; at 14 mm TL stage, no mesenchymal condensations are discerned above the endocranial cartilage (c) and the eye (e). B; the enlarged image of A for showing mesenchymal cells above the endocranial cartilage. C; the enlarged image of mesenchymal cells above the eye. Mesenchymal cells are stretched and connected via podia. D; at 17 mm TL stage, the frontal bone (f) develops dorsally to the supraorbital cartilage (c). The frontal bone is in a close proximity to a neuromast (n). E; the enlarged image of the dotted rectangle in D. The thin bone matrix (b) is surrounded by compact osteoblasts (o). F; The more lateral side of the embryos shows the initiation of the mesenchymal cell condensation for the frontal bone (f). The estimated position of the section is indicated in Figure 2. G; the enlarged image of the dotted rectangle in F. The mesenchymal cells aggregate with each other establishing a dorsoventrally single-cell layer condensation. H and I; the other sections of different embryos at 17 mm TL stage showed the different phases of the frontal bone development. The inner space for mineralization is created (arrows in H), and bone matrix is confirmed in the cell condensation (I). J; at 20 mm TL stage, the frontal bone anteriorly extends. K and L; zoomed in images of the bone above the endocranial cartilage (K) and the anterior extremity of the bone (L). Thin and stretched osteoblasts surround the extremity of the frontal bone. M; at 25 mm TL stage, the frontal bone further extends along the anteroposterior axis. N; the enlarged image shows that the osteoblasts surround the bone matrix. O; at the extremity of the bone, osteoblasts extend the podia to the bone (arrows). The inset shows the podia between the osteoblast and bone. Scales are the same in A, D, J and M (scale bar is 10 μm) and in B, C, E, F, H, I, K, L, N and O (scale bar is 100 μm). b; bone matrix, c; endocranial cartilage, e; eye, f; frontal bone, and o; osteoblasts.

At 14 mm TL, cranial mesenchymal cells were uniformly distributed between the basement membrane of epithelial cells and underlying endocranial cartilage without any signs of mesenchymal condensations for the frontal bone (Figure 3A–C). Mesenchymal cells stretched and extended long podia from the cell body, contacting each other at peripheral regions. At this stage, no signatures of osteoblast differentiation were discerned, at least at the morphological level (Figure 3B, C).

At 17 mm TL, the frontal bone started to develop dorsal to the supraorbital cartilage (Figure 3D). The bone matrix, surrounded by osteoblasts, was dorsoventrally thin (Figure 3E). Interestingly, at the more lateral position of the head, the early stage of mesenchymal condensations was discerned. Some mesenchymal cells became cuboidal and started congregating with each other dorsally to the endocranial cartilage (Figure 3F, G). Each condensation was a dorsoventral single-cell layer consisting of 2–10 cells. These cell condensations seemed to extend along the anteroposterior axis, recruiting mesenchymal cells to their extremities. In the condensations, cellular membranes of adjacent cells tightly contacted each other (Figure 3G), implying that the cells were interlinked by microscale architectures, such as gap junctions, which are indispensable for osteoblast differentiation^17,46^. Intriguingly, observing other sections in the same embryo or sections in different embryos at 17 mm TL, we could also identify slightly different stages of frontal bone development (Figure 3H, I). After mesenchymal cells started condensing, a small space was created inside the cell condensations (Figure 3H). Subsequently, bone matrix was produced into this space (Figure 3I). Contrary to a previous finding in bowfin^33^, a break of the basal membrane of the epidermis and subsequent cell migration from the neuromasts to the mesenchymal layer was not confirmed^33^.

At 20 mm TL, the frontal bone extended along the anteroposterior axis (Figure 3J). At this point, the frontal bone was a flat osseous plate surrounded by osteoblasts and grew adjacent to the supraorbital cartilage. A relatively small number of osteoblasts were observed in the middle part of the frontal bone compared to anterior and posterior extremities (Figure 3K, L).

At 25 mm TL, the frontal bone further developed and was strongly stained by alizarin red S (Figure 3M, N). The frontal bone delineated the supraorbital cartilage more than the basement membrane of epithelial cells (Figure 3M). Intriguingly, osteoblasts protruded podia to the bone matrix, implying that they communicate with the bone matrix via podia and regulate ossification (Figure 3O). 2 replicates (different individuals) were observed for each stage/observation.

### 2.4 | Cytoskeletal and membranous changes during ossification

Dermal ossification starts with the formation of mesenchymal cell condensations^47^. Multiple molecular markers for condensing cells, such as NCAM or fibronectin, were identified in previous studies^48^. However, cytoskeleton and plasma membrane changes in condensing cells remain poorly described. Therefore, we investigated F-actin (cytoskeleton) by phalloidin and the plasma membrane by membrane staining at the initiation of cell condensation. At 17 mm TL when mesenchymal cells start to aggregate for the frontal bone, 2–3 cells in the condensations showed significantly enriched F-actin staining (Figure 4A, B. n = 5). The spatial distribution of the cells with the enrichment of F-actin seems to be stochastic in the mesenchymal condensation (Figure 4B, C, D. n=5). Other cells in the condensations also displayed slightly higher F-actin signal than surrounding mesenchymal cells (Figure 4A, B). Intriguingly, aggregating cells had smaller and more condensed nuclei than surrounding mesenchymal cells (Figure 4E, I. n = 5). The DAPI staining displayed that DNA was unevenly distributed in these nuclei, implying that chromatin structure is rapidly changing, which highly likely affects the differentiation state of these cells via gene expression changes (Figure 4E, inset). Cells with the enriched F-actin signal also exhibited strong staining of the plasma membrane, indicating that the amount or composition of membrane components (e.g., phospholipids) also changes (Figure 4F-H. n = 5). Although both F-actin and plasma membrane were enriched in the same cells, the high-magnification observation showed that these two signals did not spatially overlap in the cells, implying that cytoskeletal and membranous changes may be independent or indirectly linked pathways (Figure 4H, inset).

**Figure 4.**
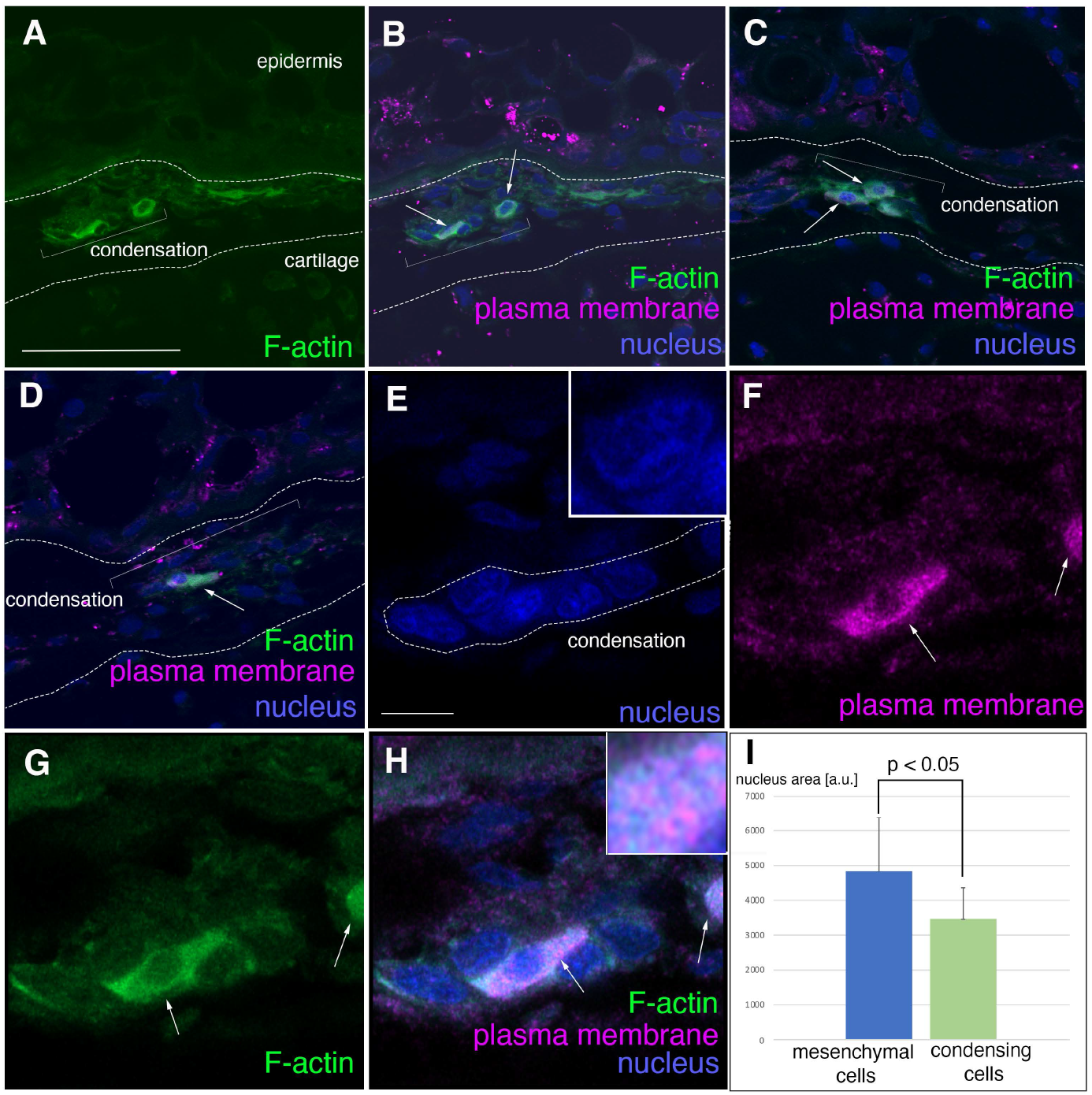
Alteration of F-actin and phospholipids organization during the cell condensation formation. A-D; F-actin staining (Phalloidin) (A), and the merged image of F-actin, phospholipids (CellMask), and nucleus (DAPI) staining (B-D) in the mesenchymal cell condensation at 17 mm TL stage. Among the condensing cells, some cells showed strong F-actin and phospholipid staining (arrowheads). B, C, and D show the sections of different individuals. Note that the localization of the cells with enriched F-actin is random in the mesenchymal condensations. E-H; high-magnification observation of the mesenchymal cell condensation. The nuclei in the aggregating cells become compact and the chromatins are unevenly distributed in the nuclei (E, inset). In the mesenchymal condensation, some random cells showed significantly higher staining of both phospholipids (F) and F-actin (G) than others. The localization of the upregulated F-actin and phospholipids do not colocalize in the cells (the inset in H). Scale bars are 100 μm in A and 10 μm in E. I; the comparison of the nuclear size (surface area) between mesenchymal cells and condensing cells (t-test, p=0.045)._

## 3 | DISCUSSION

### 3.1 | Possible mechanisms that induce mesenchymal cell condensations for the frontal bone

The position and size of each cranial dermal bone underpin the evolutionary diversity of fish skulls. As the positions of cranial dermal bones are determined by the initial mesenchymal condensations, identifying the main inducers of mesenchymal condensations is imperative to understanding fish skull diversity. Despite the critical need to uncover these mechanisms, determining the ancestral and shared processes for cranial dermal bone induction can be difficult using model organisms due to evolutionarily derived developmental characteristics^49^. Therefore, we used spotted gar as an experimental model^41^, a basally branching actinopterygian that likely retains the ancestral molecular mechanisms of fish skull roof formation. HE section staining of spotted gar juveniles showed that mesenchymal condensations developed in the uniformly distributed cranial mesenchymal cells, which were dorsoventrally flanked by the epidermis and underlying endocranial cartilage (Figure 3). The initial mesenchymal cell condensations were spatially fragmented and developed as dorsoventral single-cell layers, which are evidently isolated from the epidermis and endocranial cartilage. This finding is slightly different from the prior description of osteoblast differentiation in teleost, i.e. cichlid, frontal bone development, which showed that osteoblasts adjoin perichondral cells of the endocranial cartilage^16^. Additionally, our finding differs from the notion that epidermal cells migrate from neuromasts to the mesenchymal layer and probably form mesenchymal condensations in bowfin juveniles^33^. Given that mesenchymal condensations of the frontal bone initially develop well-isolated from the epidermis and endocranial cartilage in spotted gar, we suggest possible mechanisms to induce the mesenchymal condensations for the frontal bone below.

I. Since Wnt and Fgf ligands expressed in the cranial ectoderm (epidermal layer) regulate differentiation of osteoblasts and chondrocytes in the cranial mesenchymal layer via protein diffusion^38,50,51^, distance from the ectoderm may be a determinant of osteoblast differentiation at a certain position in the mesenchymal layer. For example, mesenchymal cells adjacent to the ectoderm may receive an excessive dose of Wnt or Fgf signal, and/or cells adjacent to the endocranial cartilage may receive an insufficient amount of signals. Expression levels of Wnts and Fgfs must be tightly regulated in vertebrate skull formation, and modification of their activities may produce the evolutionary diversity of frontal bone patterning.
II. In addition to diffusible molecules from the ectoderm to the mesenchymal layer, other diffusible molecules from the underlying meninges and cartilage, such as BMPs, regulate growth and patterning of the frontal bone condensation^50,52^. Single knockout of BMP genes in mice does not affect initial cell condensations for the frontal bone but disrupts frontal bone growth and patterning at later stages^50,52^. Thus, diffusible signals from both the epidermis and underlying cartilage may create dorsoventrally opposite signaling gradients and provide positional information to mesenchymal cells to initiate mesenchymal condensations.
III. Increase in the density of mesenchymal cells may autonomously create an uneven distribution of diffusible molecules or extracellular matrix in the cranial mesenchymal layer. As cranial mesenchymal cells proliferate, the concentration of secreted molecules from the mesenchymal cells themselves may become highest in the middle of the cranial mesenchyme layer. Intriguingly, mesenchymal stem cells are specified to osteoblasts with *Runx2* expression depending on the density of cells in in vitro collagen scaffolds^53^. Once a cell population achieves a density threshold during embryonic development, mesenchymal cells may differentiate into osteoblasts with a certain amount of diffusible molecules or extracellular matrix between the epidermis and supraorbital bone.
IV. Another possible mechanism to determine the position of mesenchymal cell condensations in the mesenchymal layer is mechanical force. A prior study in salmon (Salmonidae) suggested that mechanical force from the endocranial cartilage or the surface ectoderm may control frontal bone development^54^. Importantly, mesenchymal stem cells are reported to be specified to osteoblasts or adipocytes depending on cell shape^55,56^. Cell shape change triggers the Hippo-YAP signaling pathway^57–61^, which has been suggested to be a key player for osteoblast differentiation^61^. Although we have not tested whether cranial mesenchymal cells receive any mechanical pressure, change in the expression level of F-actin in the prospective osteoblasts may occur upon physical cell–cell interaction or shear stress^62^. This hypothesis should be carefully examined in future studies.

### 3.2 | Cell condensation forms in proximity to neuromasts, but not at the exact same positions

Previous studies in gar and bowfin have found mesenchymal cell condensations for the frontal bone under sensory organs in the epidermis^31,32,63^. Given the proximity of the cranial dermal bones to neuromast cells during development, some studies asserted that lateral line neuromasts may induce mesenchymal cell condensations for cranial dermal bones in fish^44^. Consistently, our HE-stained sections of gar juveniles indicated that neuromasts in the supraorbital line are in close proximity to mesenchymal condensations during frontal bone development. However, high resolution 3D topological analysis of neuromasts and frontal bone anlagen showed that the positions of the neuromasts and condensations do not dorsoventrally align, although they develop in a close manner. This observation is also consistent with observations in other studies in actinopterygians and crossopterygians that suggest that the close association between neuromasts and cranial dermal bones is a developmental coincidence^64,65^. Moreover, removal of neuromasts from the supraorbital line, which runs at the dorsal surface of the frontal bone, delays growth of the frontal bone without affecting initiation of frontal bone formation^66^. Collectively, neuromasts seem to be irrelevant to the induction of mesenchymal condensations for the frontal bone of spotted gar juveniles. However, close localization of neuromasts may affect frontal bone development, such as in the modification of bone growth via diffusible molecules or cell migration^33^. Intriguingly, a recent study in the caveform of the Mexican tetra *Astyanax mexicanus* discovered that ossification centers of the suborbital bones correspond with neuromast positions^67^. Also, evolutionary diversity of the suborbital bone in *Astyanax mexicanus* is correlated with neuromast distribution—this new finding may refuel the classical hypothesis or represent the evolutionarily derived mechanisms acquired in Mexican tetra. If neuromasts regulate frontal bone formation, further studies to identify the underlying mechanisms, such as cell migration from neuromasts to the mesenchymal layer^33^, would be necessary. Alternatively, the developmental relationship between dermal bones and neuromasts may be the other way around such that the developing dermal bones stimulate neuromast formation.

### 3.3 | Development of the frontal bone in spotted gar at single-cell resolution

#### I) Asymmetric information in osteoblasts for frontal bone development

While macroscopic observation of frontal bone formation in non-teleost and teleost actinopterygians has been conducted elsewhere^16,29,33,66^, description of the developmental process at the single-cell resolution is limited^16^. Our high-magnification observation of the developing frontal bone in spotted gar provides fundamental insights into the developmental process and underlying molecular mechanisms. After several mesenchymal cells aggregate, the cells start creating the inner space for bone matrix (Figure 5). This observation suggests that osteoblasts in mesenchymal condensations possess cell polarity and can establish “inside–outside” information of the condensations. In mesenchymal–epithelial transitions (MET), mesenchymal cells condense and autonomously obtain apicobasal information^68^. Although formation of mesenchymal condensations for cranial dermal bones significantly differs from MET, the osteoblast polarity information in the frontal bone anlage may be generated by similar molecular players such as Zo^68^ or other cell–cell junction proteins. Gap junctions between mesenchymal cells are indispensable for osteoblast differentiation^46^. Thus, the proteins involved in gap junctions may establish the polarity information for condensing osteoblasts. Therefore, future work should investigate the localization of cell junction molecules in the condensing mesenchymal cells in spotted gar to understand how osteoblasts develop bone matrix inward.

**Figure 5.**
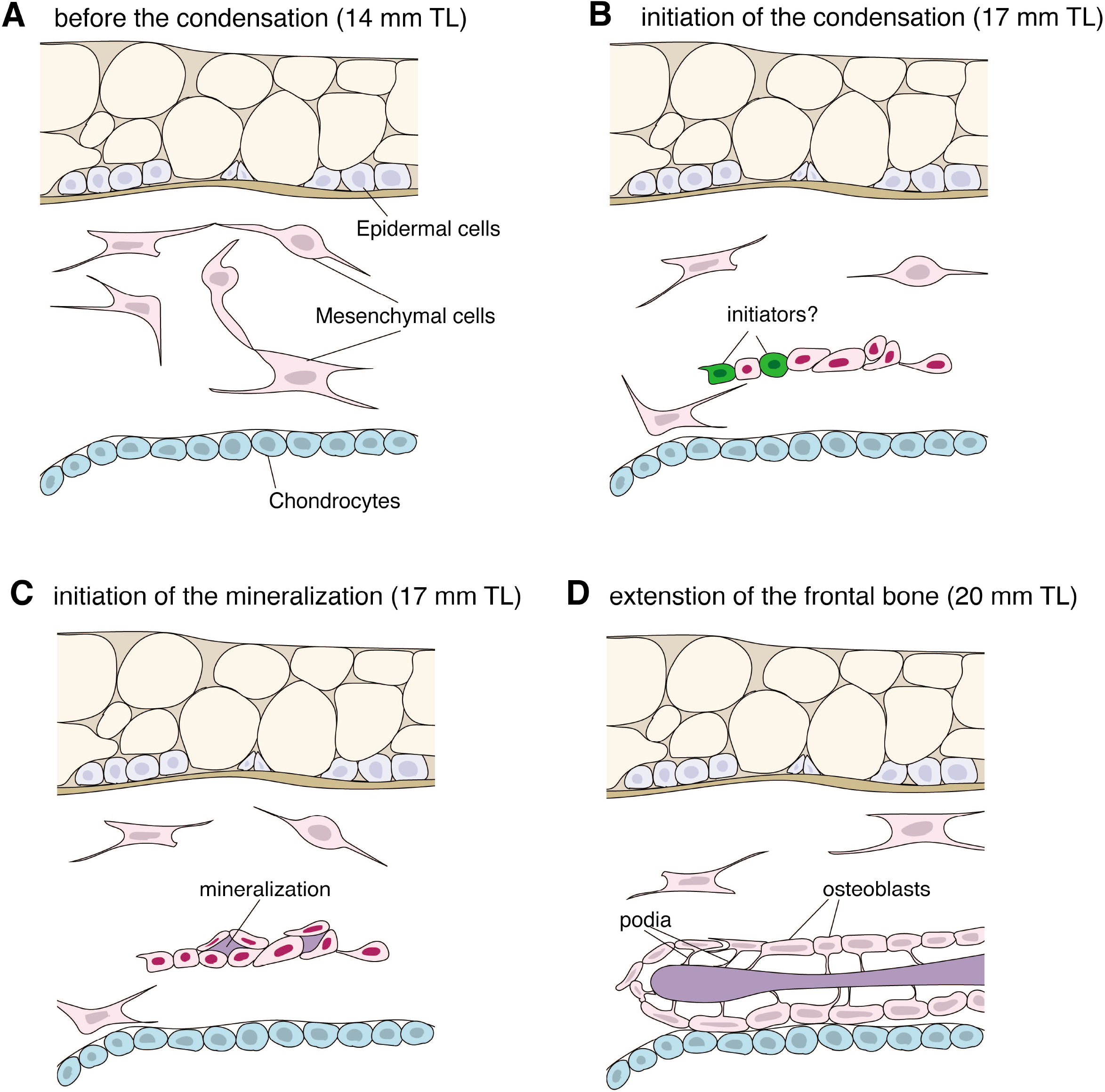
Developmental landmarks of the frontal bone in spotted gar. A; before the formation of the mesenchymal condensations. The mesenchymal cells stretch and protrude multiple podia from the cell bodies. B; when the mesenchymal cells aggregate, a couple of random cells in the condensation display the upregulation of F-actin and phospholipids (green). These cells may promote the condensation process or stimulate osteoblast differentiation. C; The mesenchymal condensations produce bone matrix inside. The osteoblasts may have “inside-outside” positional information. D; as the frontal bone grows, osteoblasts protrude podia to the bone matrix. These podia may supply calcium or other minerals to the bone matrix and regulate bone growth.

#### II) Osteoblasts articulate to the bone via podia

After mesenchymal condensations are formed, the mineralized frontal bone extends along the anteroposterior axis. Intriguingly, we found that osteoblasts protrude podia to the bone matrix at extremities of the developing frontal bone (Figure 5D). A similar structure was briefly observed in osteoblasts in a cortical bone of the mouse tibia^69^, yet the detailed structure remains undescribed. Since all osteoblasts on the surface of the bone matrix possess podia connecting to the bone matrix, these podia potentially have critical functions in bone development, such as supplying phosphate or calcium, one of the main functions of osteoblasts during ossification^70^. Further, the polarity information from osteoblasts at the early stage may be carried over to later stages to instruct podia protrusion to the bone matrix. Further studies characterizing the structure of these podia using molecular staining for cytoskeletal proteins with high-resolution microscopy will promote understanding of this unique architecture and function.

### 3.4 | Cytoskeletal and membranous changes in cell condensations

Despite accumulating knowledge regarding molecular markers for the developing mesenchymal condensation^47^, the mechanisms that initiate mesenchymal cell condensations remain elusive. We stained the F-actin filaments and plasma membrane to monitor how the cytoskeleton and membrane architecture change during condensation development. Intriguingly, a couple of cells in the initial condensations showed high F-actin and plasma membrane staining (Figure 4). In cell–cell adhesion, intercellular proteins involved in cell–cell junctions (e.g., gap junctions or tight junctions) play major roles^71^. Thus, upregulation of F-actin may promote cell– cell adhesion among condensing mesenchymal cells via junction proteins. The stochastic upregulation of F-actin in a small subset of cells in mesenchymal condensations may support the existence of “initiators” that promote cell condensation (Figure 5B). Alternatively, upregulation of F-actin may occur in all condensing cells in a temporal manner but at different time points; our staining method may not have enough temporal resolution to detect this event.

Another possibility is that upregulation of F-actin controls osteoblast differentiation. F-actin plays a fundamental role in the differentiation of mesenchymal stem cells to osteoblasts via the Hippo-YAP pathway in vitro^72^. Thus, upregulation of F-actin in stochastic cells in spotted gar skull formation may be critical for osteoblast differentiation via cell–cell adhesion, which, in turn, induces osteoblast differentiation, potentially via the Hippo-YAP pathway.

Intriguingly, our results showed that the amount of plasma membrane was upregulated in the same cells with high F-actin signal. A previous study in rats showed that changes in membrane lipid composition control cell–cell adherence potency via regulation of junction proteins^73^. Thus, upregulation of the membrane phospholipid in mesenchymal condensations may promote cell–cell adhesion in parallel with upregulation of F-actin. We found that F-actin and phospholipid localization do not completely overlap in condensing mesenchymal cells, suggesting that upregulation of F-actin and phospholipids probably regulates cell condensation formation in an independent and cooperative manner.

### 3.5 | Ancestral and conserved mechanisms of frontal bone development in basal actinopterygians

We identified initiation and growth processes of the frontal bone in spotted gar at single-cell resolution. Although the patterning of cranial dermal bones, including the frontal bone, diversified over 400 million years, the central facet of skull development seems to be conserved with myriad minor modifications to adapt to distinct habitats and functional necessity. Given the phylogenetically prominent position of gar in vertebrate evolution, the newly identified mechanisms, such as the upregulation of F-actin and plasma membrane or the podia from osteoblasts to the frontal bone, are likely conserved features of cranial bone development in other fish. Further research should test the conservation of these findings in other actinopterygian and teleost fish. Future studies to identify the upstream regulators of cytoskeleton and membrane changes, along with their functions, would illuminate fundamental mechanisms of cranial dermal bone development and evolution in vertebrates.

## 4 | EXPERIMENTAL PROCEDURES

### 4.1| Animals

Animal work was approved under Institutional Animal Care and Use Committee (IACUC) protocol #10/16-179-00 from Michigan State University. Spotted gar from the Louisiana population were spawned and raised as described previously (citation: PMID 24486528) with a few modifications. Embryos and larvae were raised up to the desired stages in plastic containers of increasing size in culture water made from RO water and Instant Ocean salt, at 18-20 degree Celsius, and with a light cycle of 12 hrs light/12 hrs dark. Later stages were fed artemia nauplii at least twice a day. At the target stages, gar embryos were euthanized by an overdose of tricaine and fixed in 4 % PFA, 10 % formalin, or Bouin’s fixative at 4 degree Celsius overnight. The total length (TL) of each specimen was measured using a stereotype microscope (Leica M250) and associated software.

### 4.2 | Acid-free Bone and cartilage staining

Bone and cartilage staining without acetic acid was performed as previously described^42^. Prior to staining, specimens fixed in 10 % formalin were briefly rinsed in distilled H_2_O (dH_2_O) twice. The entrails of specimens were removed for large specimens by tweezers or a small slit was carefully made at the ventral side of the body for small specimens in dH_2_O. Then, specimens were rinsed in dH_2_O on a rocker at room temperature overnight. The next day, specimens were equilibrated in 70 % ethanol solution and cartilages were stained in Alcian Blue 8 GX in 70 % ethanol / 50 mM MgCl2 overnight. After the staining, specimens were equilibrated to dH_2_O through an ethanol: dH_2_O series (3:1, 1:1, 1:3) and immersed in a 1 % trypsin: 30 % saturated sodium borate solution until specimens became translucent. Subsequently, bones were stained in alizarin red solution (0.005 % Alizarin Red S powder in 1 % potassium hydroxide) on a shaker overnight. After bone staining, the specimens were bleached for one week in a solution (75 % of 0.1 % potassium hydroxide in water and 25 % of glycerol). Specimens were transferred to 100 % glycerol through a 1 % potassium hydroxide: glycerol series (1:1, 1:3).

### 4.3 | Paraffin section and HE staining

Bouin-fixed gar embryos were subjected to paraffin sectioning and HE staining. Longitudinal sections (8 μm) were made at 14 mm, 17 mm, 20 mm, 22 mm, and 25 mm TL stages by the Research Pathology Services at Rutgers University. The slides with mounted sections were then stained by Hematoxylin and Eosin solutions. The stained sections were enclosed by cover-glasses and then photographed by an Olympus BX63 (Imaging Core, Human Genetics Institute of New Jersey, Rutgers).

### 4.4 | 3D reconstruction of bone development in gar embryos

The images of 35 serial HE stained sections at 17 mm, 20 mm and 25 mm TL stages were incorporated into 3D visualization and analysis software Amira (Thermo Fisher) and the positions of all sections were manually aligned. Then, the eye, endocranial cartilage, neuromasts, and frontal bone were manually segmented out, reconstructed to 3D, and pseudo-colored.

### 4.5 | Cell membrane and actin cytoskeletal staining

Gar embryos fixed in 4 % PFA were soaked in a series of sucrose solutions (10 %, 15 %, 20 %) and embedded in OCT compound (Sakura tissue). Then, the OCT blocks were flash-frozen in liquid nitrogen. The cryosections were made at 8 um thickness and mounted on slide glasses. Upon the removal of OCT compound in PBS with 0.1 % Triton X-100 (PBT), the sections were stained by Alexa Fluor™ 488 Phalloidin (1/1000 dilution, ThermoFisher Scientific), CellMask (1/1000 dilution, ThermoFisher Scientific), and with DAPI (1/4000 dilution). After three brief washes in PBT, sections were covered by cover glasses with 50 % Glycerol and dH_2_O. Then, the staining signals for F-actin, phospholipids, and nuclei were photographed using a confocal microscope (Zeiss LSM 510).

### 4.6 | Quantification of nuclear size in mesenchymal cell condensations

The confocal scanned images of mesenchymal condensations stained by F-actin, CellMask, and DAPI were loaded on ImageJ and surface are of nuclei was measured. For condensing cells and surrounding mesenchymal cells, 8 or 10 nuclei were quantified, respectively.

## ACKNOWLEDGEMENTS

This work was performed with the institutional support provided by the Rutgers University School of Arts and Sciences and the Human Genetics Institute of New Jersey (to T.N). We thank Solomon David and Allyse Ferrara (Nicholls State University) for their help with collecting gar broodstock and eggs as well as Camilla Peabody (Michigan State University) for help with gar embryo husbandry.

## Supplementary Information

**Supplementary Figure 1.**
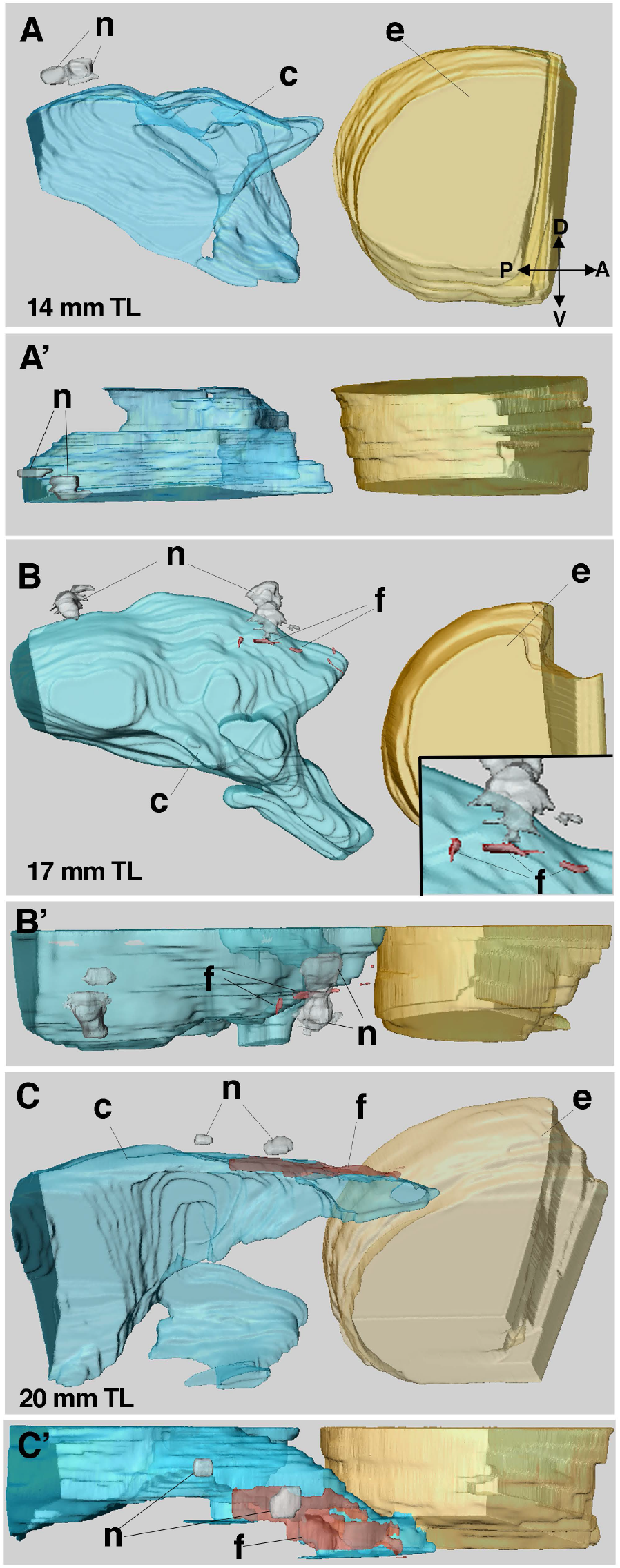
The topological relationship among the neuromasts, endocranial cartilage, and frontal bone at the peripheral side of the juveniles. A, B, and C; the dorsolateral views of 3D reconstruction at 14 mm (A), 17 mm (B), and 20 mm (C) TL stages. A’, B’, and C’; the dorsal views of the same juveniles of A, B, and C. At 14 mm TL stage, the frontal bone anlage is not observed (A and A’). At 17 mm TL stage, compared to the developing frontal bone at the medial part of the juveniles (Figure 2), the several mesenchymal cells start to aggregate and produce the mesenchymal condensations (f) at the peripheral side (B). The condensation occurs in a close proximity to the neuromasts (n), but condensations are located between two neuromasts in the mesenchymal layer (B’). At 20 mm TL stage, the frontal bone at the peripheral side starts to expand anteriorly and posteriorly (C and C’). While the frontal bone forms under the neuromasts, it anteriorly and laterally expands without neuromasts (C’). The scale bars are 0.2 mm. c; endocranial cartilage, e; eye, f; frontal bone, and n; neuromasts.

